# Eye-Tracking-BIDS: the Brain Imaging Data Structure extended to gaze position and pupil data

**DOI:** 10.64898/2026.02.03.703514

**Authors:** Martin Szinte, Dominik R. Bach, Dejan Draschkow, Oscar Esteban, Benjamin Gagl, Rémi Gau, Klara Gregorova, Yaroslav O. Halchenko, Scott Huberty, Sina M. Kling, Sourav Kulkarni, The BIDS Maintainers, Christopher J. Markiewicz, Mark Mikkelsen, Robert Oostenveld, Julia-Katharina Pfarr

## Abstract

The Brain Imaging Data Structure (BIDS) is a widely adopted, community-driven standard to organize neuroimaging data and metadata. Although numerous extensions have been developed to incrementally extend coverage to new modalities and data types, an unambiguous, granular specification for eye-tracking recordings is lacking. Here, we present how BIDS will structure data and metadata produced by eye-tracking devices, including gaze position and pupil data. In addition to prescribing the organization of the unprocessed (raw) recordings and associated metadata as produced by the device, BEP20 also resolves gaps in current BIDS specifications beyond the scope of eye tracking. In particular, it adds a mechanism for including asynchronous model parameters and messages, such as contextual information, statuses, and events, such as triggers, generated by the device. BEP20 includes examples that illustrate its applicability in various experimental settings. This BIDS extension provides a robust standard that supports the development of self-adaptive, open, and automated eye-tracking data structures, thereby bolstering transparency and reliability of results in this field.

## Introduction

Advancements in modern eye-tracking technologies allow researchers and developers to track eye movements and pupil size^1^. At the heart of eye-tracking lies the capability to assess the gaze position, typically achieved through specialized cameras and software. Indeed, most contemporary eye-trackers are video-based, utilizing infrared light to create a reflection on the cornea (tip of the eyeball) that can be tracked by the system. By calibrating the eye-tracker to the user’s gaze, the software can accurately determine where an individual is looking on a screen or in the real world with high spatial and temporal precision. Moreover, some technologies now allow the use of simple webcams or of embedded eye-trackers within virtual and augmented reality sets. These recent advances promise to expand access to gaze and pupil data, enlarging an already well-developed community of users.

Eye movements allow the human eye to scan a scene, providing the visual system with the relevant information to understand our environment^2,3^. Various types of eye movements, such as voluntary saccades, smooth pursuit, or involuntary nystagmus, enable the high-acuity region of the retina (the fovea) to extract detailed information from different parts of static or moving visual scenes^3^. Interestingly, gaze data offer insights into cognitive processes such as language comprehension^4,5^, memory^6,7^, attention^8,9^, mental imagery^10^, and decision-making^11,12^. Moreover, most eye-trackers also provide information about pupil size, which has been found to relate to different cognitive variables in emotion research^13^, decision-making^14^, and associative learning^15^. Pupil size is controlled by the locus coeruleus and might provide direct information on this brain region’s activity, underlining the potential importance of this data type^16^. While the use of eye-tracking data varies across domains and scientific questions, it is often collected alongside other observables, such as behavioral responses or neural activity (e.g.,EEG, MEG, fMRI, and electrophysiology), thus contributing to a rich understanding of the studied cognitive processes.

Sharing research data, including eye-tracking data, is valuable and often mandatory for various reasons. First, it is considered good scientific practice to share the results of published research in accordance with the FAIR principles so that others can reproduce the results with or without the same research data^17,18^. Second, there are increasing incentives to reuse existing data to generate new research questions. To this end, many funders require that original research data be deposited for others to use as well as to maximize the impact of researchers’ contribution. Third, by collecting and combining large samples of original data, mega-analyses of existing research can overcome the shortcomings associated with underpowered studies in smaller samples^19^ and thus complement the traditional meta-analytic approach of combining statistical results. However, in practice, there are significant barriers to using shared data when it is deposited in multiple formats, as aligning these formats consumes resources. Indeed, the diversity of eye-tracking apparatus and their users results in a non-standardized variety of data and metadata, often relying on proprietary eye-tracker formats such as EyeLink’s *Eyelink Data Format* (EDF, not to be confused with the European Data Format^20^, an EEG file format employing the same file name extension). Despite efforts to develop good practices for organizing eye-tracking data^21,22^, and related standardization efforts, such as Neurodata Without Borders (NWB)^23^, which includes eye-tracking data within its behavior module, researchers continue to develop their own, idiosyncratic data-management protocols, which largely limit data sharing and reuse. Thus, a community standard for domain-specific data types could help put the general notion of open data into actionable practice.

To this end, BIDS^18^ emerged as a powerful standard for describing and sharing data in the neuroimaging research community. While BIDS initially started with only one specification for MRI data annotation and organization, it has been continuously extended to other recording modalities (for a retrospective^24^) such as EEG^25^, MEG^26^, PET^27^, iEEG^28^, fNIRS^29^, and motion tracking^30^. Our approach facilitates bridging existing tools, such as MNE-Python^31^ for data analysis and PsychoPy^32^ for stimulus presentation, within BIDS. While BIDS already allows the representation of most forms of eye-tracking data and metadata, by 2026 it still lacked detailed specifications for saving gaze position and pupil information that provide the necessary comprehensiveness and accuracy for reproducible and effective data sharing. By adhering to the FAIR principles of findability, accessibility, interoperability, and reusability^17^, BIDS—now including eye-tracking data—promotes data sharing, reproducibility, and automated analysis workflows. Eye-Tracking-BIDS, as it provides a structured framework for storing and describing data recordings obtained with eye-tracking devices, regardless of the specific recording technology or settings. The new specification will bolster open science practices in the eye-tracking community, facilitating data sharing, reducing replication issues, and encouraging the development of automated data analysis tools.

### Addressing the currently minimal coverage of eye-tracking within BIDS

BIDS datasets are organized hierarchically in subject-specific subdirectories (e.g., *sub-01/, sub-02/*) and, if applicable, in session-specific subdirectories (e.g., *ses-01/, ses-02/*). Within these folders lie modality-specific directories (e.g., *beh/* for behavioral data, *func/* for fMRI data, etc.). Data, in a variety of formats such as NIfTI, tab-separated values (TSV), compressed TSV, etc., and metadata, in JavaScript Object Notation (JSON), are then stored in modality-specific folders following predesignated naming conventions. In general, names are composed of “entities” which are key–value pairs separated by ASCII underscore characters, such as *sub-A01_ses-01_\**.*nii*.*gz*, to indicate a particular subject and session. Some of these entities are mandatory, whereas others are optional, as explicitly described in the BIDS specification. A special entity, the “suffix”, is placed at the end of the file to indicate specific data types such as *T1w, T2w, bold, dwi, eeg*, or *physio*. In addition to the suffix and subject entities (as well as the session entity for multi-session datasets), some datatypes require additional mandatory entities, for example, “task-” in the case of functional and behavioral data. In addition, BIDS prescribes the encoding of other relevant data in neuroimaging experiments, such as “events” files and asynchronous descriptions of experiment timings. Finally, BIDS encodes dataset-level metadata with files at the root directory. First, BIDS datasets include a file that provides an overview of the study, including information about authors and acknowledgments (*dataset_description*.*json*) and another that lists all study participants alongside optional demographic information such as age, gender, or handedness (*participants*.*tsv*).

Before this extension proposal, the specification included generic mechanisms for encoding physiological recordings (including eye-tracking) using the “*physio-*” prefix, optionally with the “*recording-*” entity. When gaze position and other eye-tracking recordings, such as pupil diameter or area, are associated with, for example, fMRI or EEG, data will be stored in their corresponding modality folder (*func/* or *eeg/*, respectively). If eye-tracking is the only modality of the experiment, data can be stored in the behavioral folder (i.e., *beh/*). As with any physiological recordings, eye-tracking data must be stored as a continuous, uniformly sampled recording as a gzip-compressed TSV file (TSVGZ). These TSVGZ files must be accompanied by a sidecar JSON file defining the columns in the file, *“SamplingFrequency*” and “*StartTime*”, which are necessary to reconstruct the reference timeframe.

Eye-Tracking-BIDS addresses several underspecifications that made eye-tracking encoding generally ambiguous and insufficient for some metadata. This extension builds on a new metadata field, “*PhysioType*”, that permits the prescription of mandatory and recommended metadata fields specific to the “*physio*” files that correspond to a given JSON file. When *PhysioType* is set to *“eyetrack*”, the specification unfolds specific metadata fields relevant to eye-tracking, such as *RecordedEye*. An additional problem this extension addresses is the need for more tabular formats beyond the *events* files to describe asynchronous timing information and annotations of experiments, such as status messages issued by the device (e.g., reporting the outcomes of the eye-tracker calibration protocol) and event annotations (e.g., recording start, pause, and stop events). While coverage of these metadata was comprehensive for the most mature data modalities in BIDS, such as MRI, it was particularly limited for physiological recordings. Eye-Tracking-BIDS introduces *“physioevents*” tabular files (in TSVGZ format) to encode metadata logged by the eye-tracker and asynchronous device-generated recordings, such as saccades or blinks.

### Encoding eye-tracking data in BIDS

Let us consider an fMRI experiment collected in a single session with a binocular eye-tracker during a visual search task as an example (Figure 2). BIDS encodes this experiment with the entity “*task-visualSearch*”, which is included in each participant’s files. In addition to the fMRI data and metadata (in black), we now include raw eye-tracking data files (in green) and corresponding metadata files (in yellow) using the “*physio*” suffix. We can also include asynchronous data and metadata with the corresponding “*physioevents*” files, including tabular data (in red) and sidecar JSON files (in brown). Because the eye-tracker is binocular, the specification recommends separating the data from each eye using the “*recording*” entity, producing two sets of files labeled with *recording-eye1* and *recording-eye2*.

**Figure 1.**
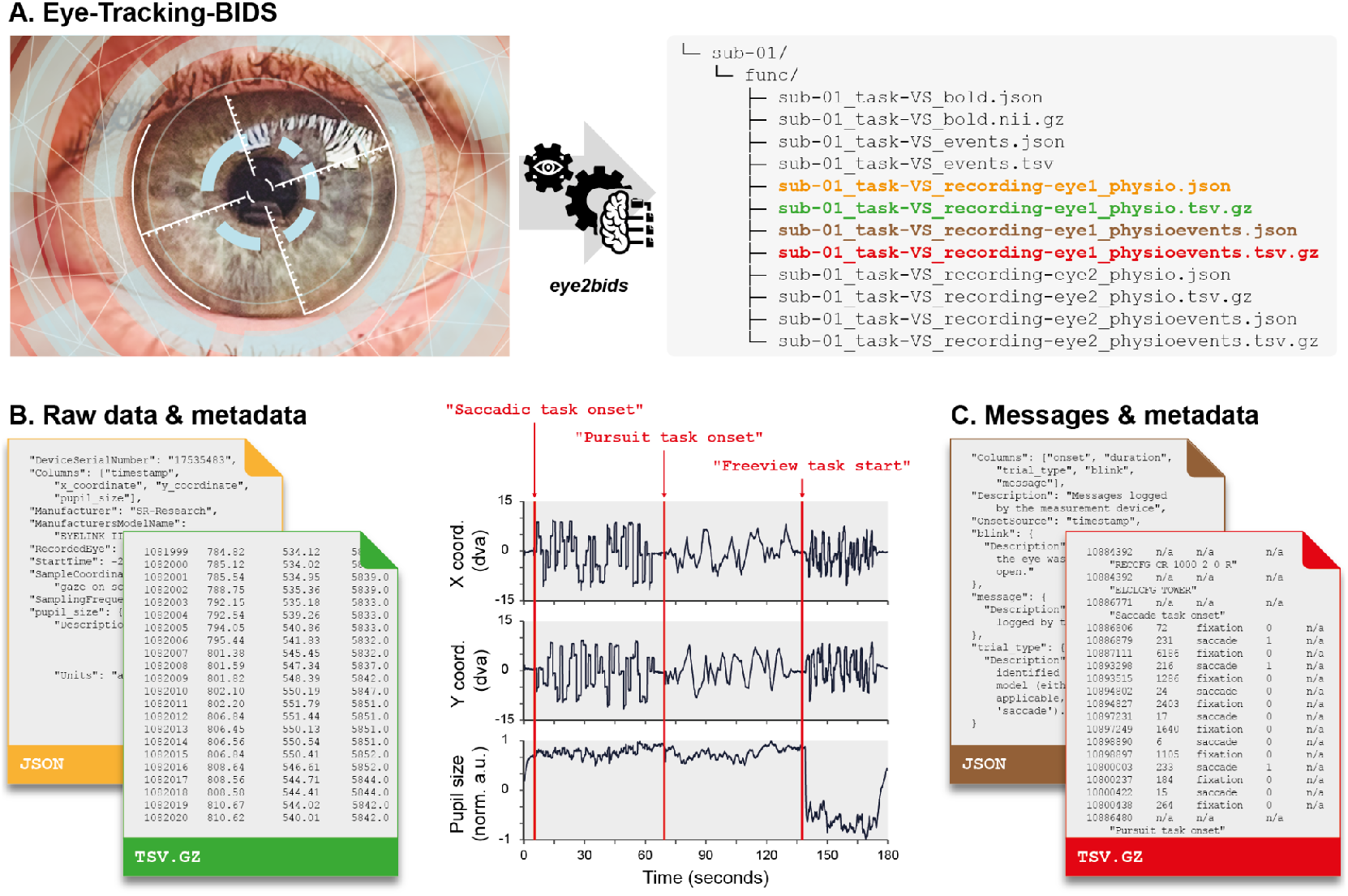
A. Eye-Tracking-BIDS. Raw data produced by the eye-tracker are organized in BIDS using the “*eye2bids*” (https://github.com/bids-standard/eye2bids) converter tool. Here, we present the typical organization of a BIDS dataset for a single participant’s binocular eye-tracking recordings combined with fMRI. **B. Storing eye-tracking raw recordings and metadata**. Gaze (x and y coordinates) and pupil timeseries plotted from data and metadata files. **C. Storing messages and metadata**. Device-generated messages are saved in a separate file, along with the corresponding metadata in two other files.

**Figure 2.**
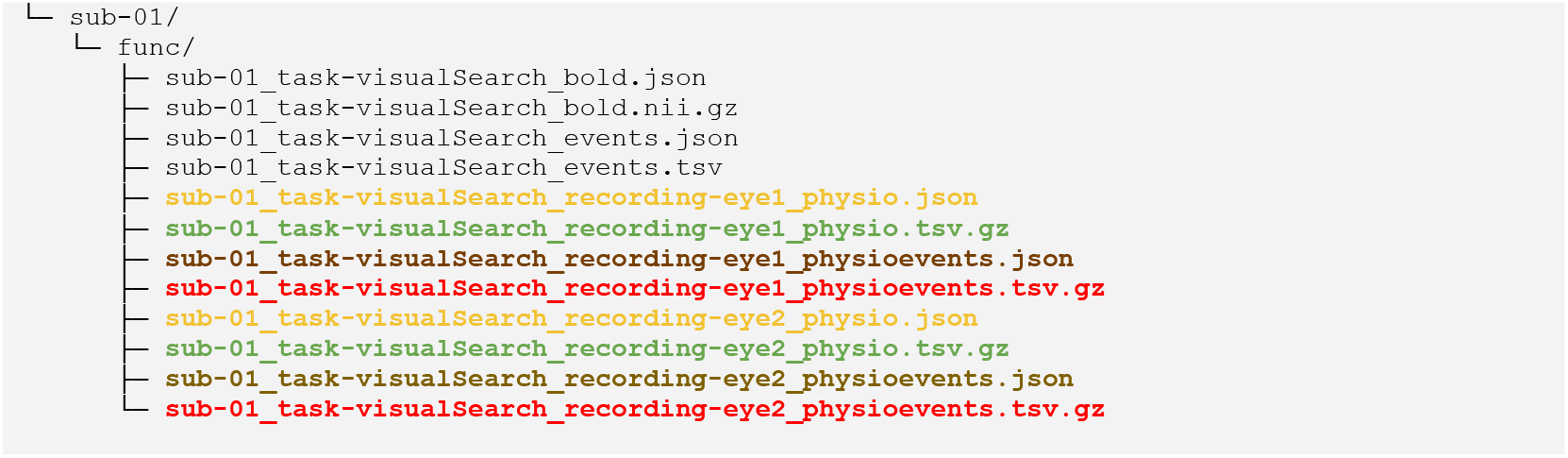
Example Eye-Tracking-BIDS directory structure of an experiment with binocular eye-tracking data.

#### Encoding metadata

Sidecar JSON files (in yellow in Figure 2) contain relevant metadata. An example of metadata encoding, corresponding to the file structure of Figure 2, is shown in Figure 3. Eye-tracking recordings continue to follow the general BIDS specifications as continuous recordings. Therefore, a “*SamplingFrequency*” with a numerical value in Hertz (e.g., 1,000) must always be defined, and a “*StartTime*” in seconds referenced to the start of the acquisition of the first data sample in the corresponding dataset (e.g., the collection of fMRI during the visual search task in our example above). If the acquisition precedes the main modality recording (e.g., fMRI), “StartTime” should have a negative value. Similarly, a “*Columns*” entry with a list of column names in the same order as their encoding in the corresponding TSVGZ file must be present. Finally, column names may be used as metadata field names, such as “*pupil_size*” in the example below, with JSON objects to refine that column’s definition, stating, for example, a “*Description*” or “*Units*”. Other fields, such as “Manufacturer” or “ManufacturersModelName,” are optional but recommended. By setting a “*PhysioType*” field to the reserved keyword “*eyetrack*”, several metadata entries become mandatory or recommended. In particular, the “recording” entity in the filename and the “*RecordedEye*” metadata in the sidecar JSON file both become mandatory. “*RecordedEye*” must take one value of “*left*”, “*right*”, or “*cyclopean*” (in the case of binocular eye-trackers that produced a third data stream with average information from both eyes’ recordings). Correspondingly, the associated \**_physio*.*tsv*.*gz* files must contain recordings for at most one eye, and the specification recommends using indexed names for the recording entity (e.g., eye1 and eye2). One additional metadata entry that becomes mandatory is “*SampleCoordinateSystem*”, which describes the eye-tracker’s coordinate system. Mandatory, recommended, and optional metadata are described in the specification under the section about physiological recordings (https://bids-specification.readthedocs.io/en/latest/modality-specific-files/physiological-recordings). As for BIDS in general, users are recommended to include any relevant settings and metadata in JSON sidecar files, even if they are not explicitly mentioned in the specifications.

**Figure 3.**
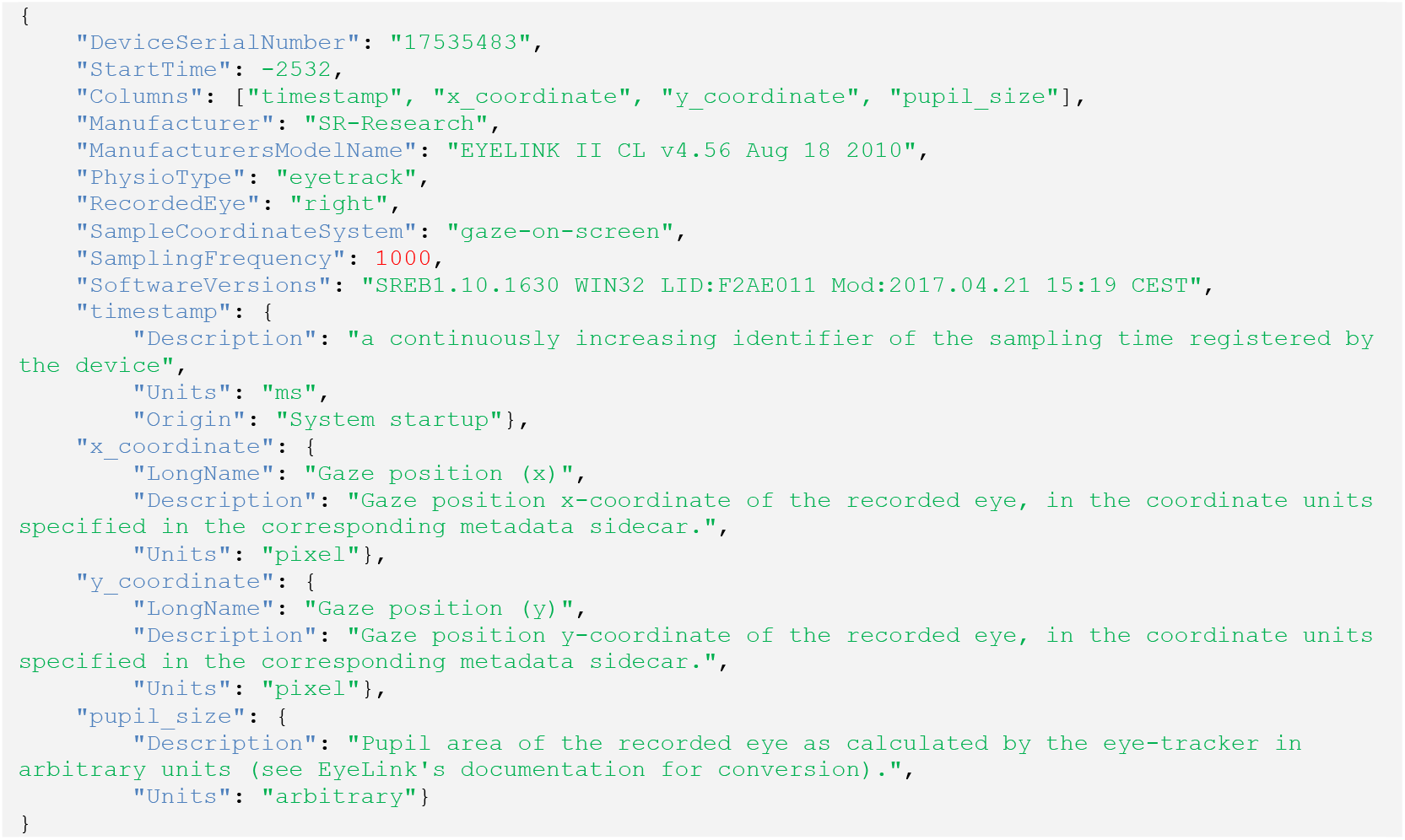
Example Eye-Tracking-BIDS metadata file (e.g. sub-01_task-visualSearch_recording-eye1_physio.json).

#### Encoding eye-tracking recordings

The raw (in the sense of device-produced) eye-tracking recordings (green in Figure 2) are stored in TSVGZ format (Figure 4) for continuous recordings. These tabular files must have at minimum three columns: a timestamp (which must appear first), and *x_coordinate* and *y_coordinate* positions of the tracked gaze with respect to the coordinate system defined by the relevant metadata entries (second and third columns, respectively). In addition to these three columns, a *pupil_size* column is optional, and the user may add arbitrary columns as long as they are BIDS-compliant (i.e., they are completely described by metadata entries in the corresponding sidecar JSON).

**Figure 4.**
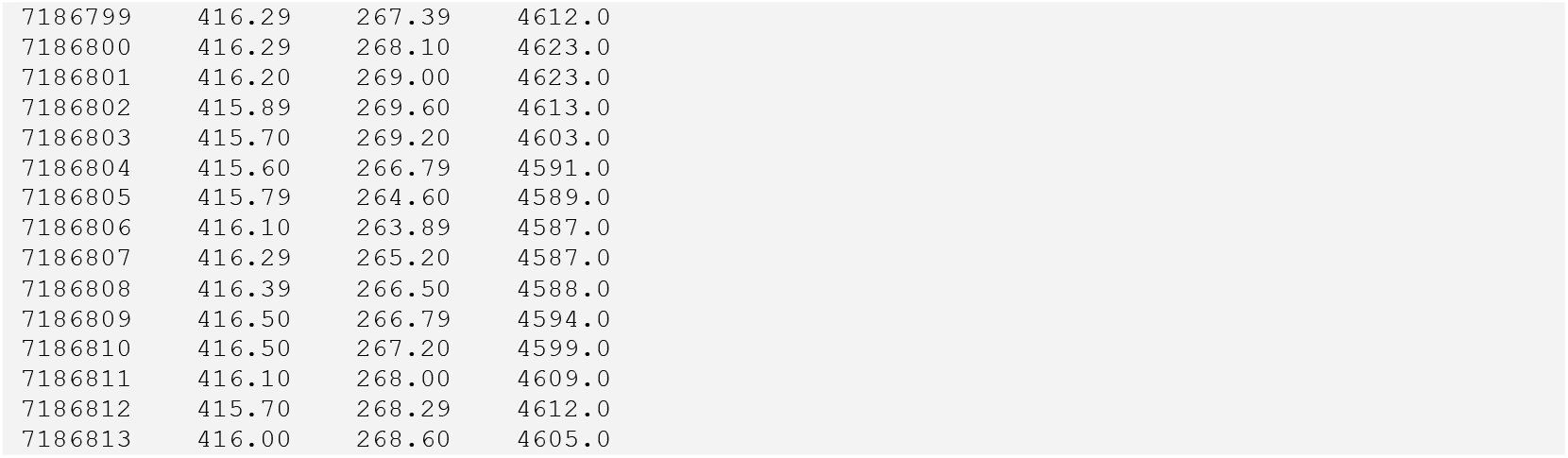
Example contents of an eye-tracker recording (e.g., sub-01_task-visualSearch_recording-eye1_physio.tsv.gz).

#### The new “*physioevents*” file

Next, we define asynchronous data files (red in Figure 2) that contain discontinuous device-generated messages or eye-tracker models and derived parameters (e.g., saccades or blinks), which are saved alongside the raw data (see Figure 5). These events are saved as single lines, each referencing the timestamps stored in the raw data file’s user-defined columns.

**Figure 5.**
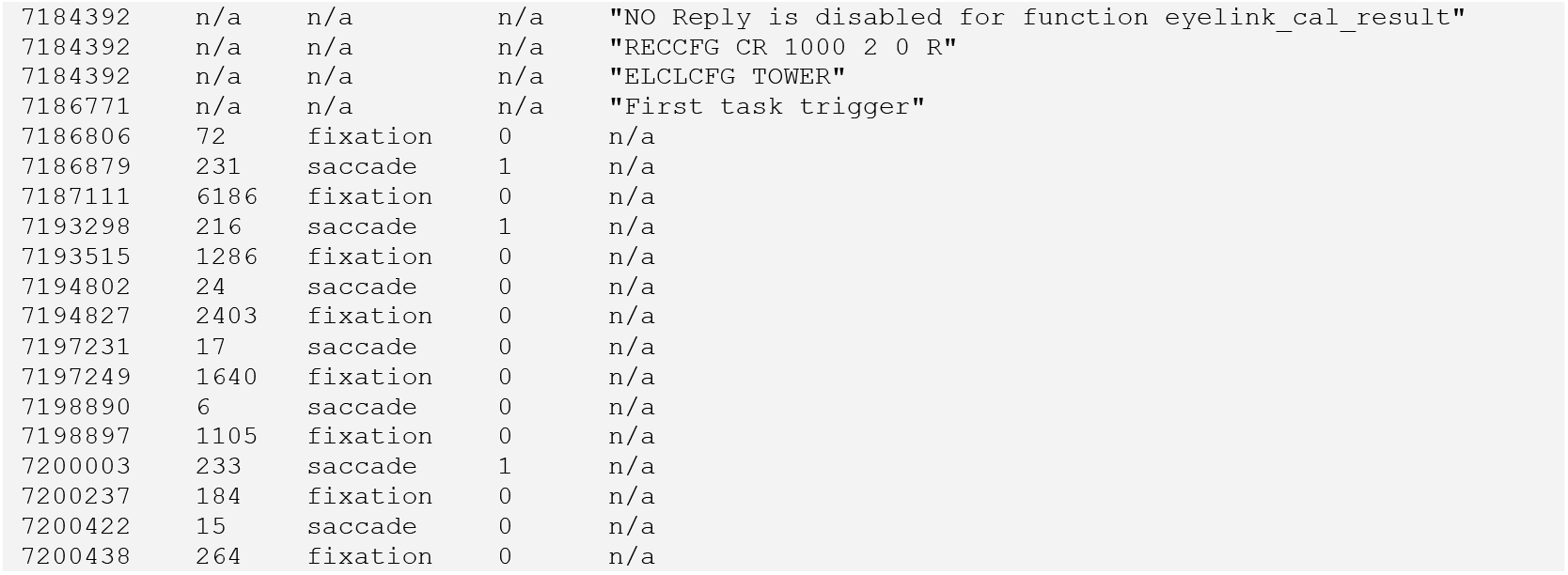
Example contents of a “*physioevents*” file (e.g., sub-01_task-visualSearch_recording-eye1_physioevents.tsv.gz).

The corresponding metadata file (in brown in Figure 2) describes the user-defined columns and the column to which the events refer, “*OnsetSource*” (see Figure 6).

**Figure 6.**
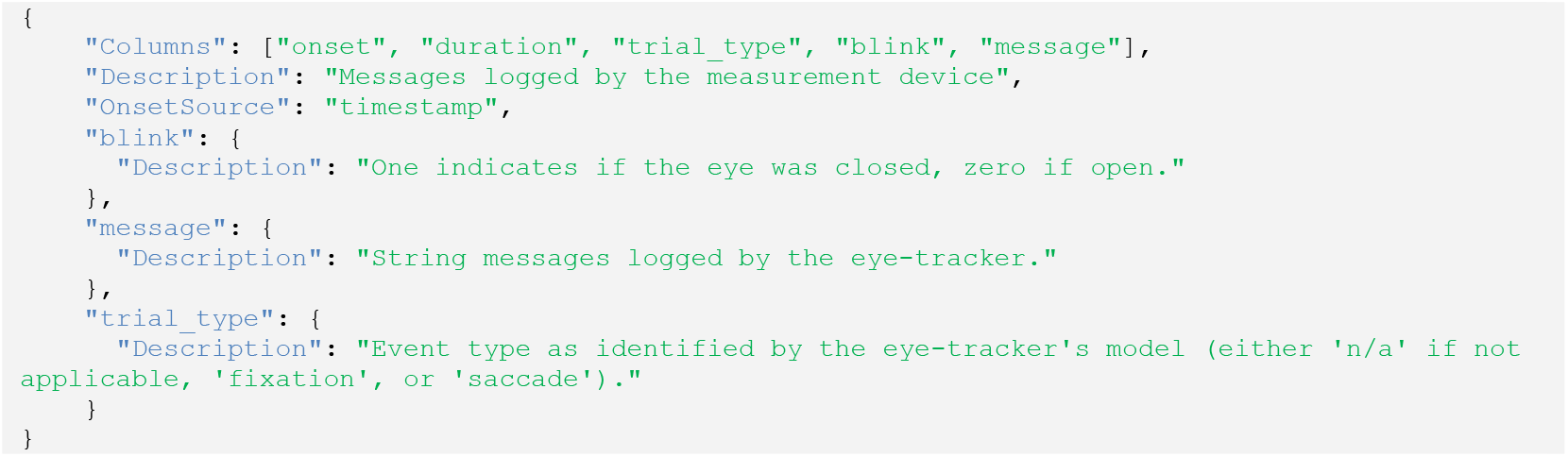
Example contents of a “*physioevents*” file (e.g., sub-01_task-visualSearch_recording-eye1_physioevents.json).

### Conversion tools and simple example datasets

To demonstrate the feasibility and effectiveness of this extension to encode eye-tracking data, we developed a conversion tool called “eye2bids” (https://github.com/bids-standard/eye2bids). Eye2bids retrieves information from the data files produced by eye-tracking devices and generates the corresponding encoding according to this proposal. As of today, the tool handles the conversion of data generated by one of the most widely used eye-trackers (SR Research Eyelink). Future community development will seek to widen eye2bids’ support for other eye-trackers. Importantly, eye2bids ensures that the converted dataset is compliant with established BIDS standards and passes the validation check by the BIDS Validator^33^ (RRID:SCR_017255)(https://github.com/bids-standard/bids-validator). The proposal includes a set of representative example datasets. The first example shows the organization of a multimodal experiment including monocular gaze position and pupil size recordings within a resting-state functional MRI experiment (https://github.com/bids-standard/bids-examples/tree/master/eyetracking_fmri). The second example demonstrates a behavioral dataset with a free-viewing task using binocular gaze position and pupil size records (https://github.com/bids-standard/bids-examples/tree/master/eyetracking_binocular). Please note that full documentation, including more examples, can be found on the BIDS website (https://bids-specification.readthedocs.io/). The description of eye tracking data as described here is part of the BIDS specification version 1.11.0.

## Conclusion

Eye-tracking has advanced technologically, gaining attention for its potential to reveal unprecedented insight into cognitive processes. However, the diversity of eye-tracking technologies, combined with disparate data management and processing practices, hinders effective data sharing and scientific transparency. The wide and fast adoption of BIDS, which has rapidly standardized how neuroimaging data is organized, motivates the extension of the standard to precisely and uniformly describe eye-tracking experiments while also adhering to the FAIR principles. In addition to extending BIDS’ coverage to eye-tracking, this proposal introduces a more general file (the “*physioevents*” file) to formalize temporal information such as logging and other device-generated annotations. Adoption of Eye-Tracking-BIDS is expected to make eye-tracking datasets more interoperable and shareable, strengthen reproducibility, and enable automated quality control, preprocessing, and analysis pipelines akin to BIDS Apps^34^. Collectively, this standardization will catalyze collaboration and accelerate scientific progress in eye-tracking research.

## Acknowledgments

We are grateful to the members of the INVIBE team in the Institut des Neurosciences de la Timone for helpful comments and discussions, and to Alice Szinte, Clémence Szinte, and Candice Duval for their support. We thank all researchers who shared their data with us and helped develop and test the converter tool.

## Code Availability

The code for the aforementioned eye2bids conversion tool is available online under the MIT license (https://github.com/bids-standard/eye2bids). Additionally, two example dataset structures are available: https://github.com/bids-standard/bids-examples/tree/master/eyetracking_fmri, https://github.com/bids-standard/bids-examples/tree/master/eyetracking_binocular.

